# EMAlign: accurate alignment of cryo-EM maps through main-chain probability using deep learning

**DOI:** 10.64898/2026.06.21.733642

**Authors:** Hong Cao, Ji Chen, Tao Li, Sheng-You Huang

## Abstract

Accurate alignment of cryo-EM density maps is essential for comparing conformational states, searching map libraries, and guiding atomic model building, but remains challenging for noisy experimental maps and partially overlapping structures. Existing alignment methods are often based on raw maps, which may result in reduced accuracy due to the density noise, or require manual intervention for local alignment, which suffers from limited general applicability. Addressing the limitations, we present EMAlign, an automatic global and local cryo-EM map alignment with predicted main-chain probability using deep learning. First, EMAlign predicts main-chain probability maps from raw cryo-EM density maps using a BiMCUNet network. Then, a fast Fourier transform (FFT)-based search strategy is used to globally search the accurate alignment between cryo-EM maps based on predicted main-chain probability maps. As such, the main-chain probability map overcomes the noisy raw map problem, and the FFT-based exhaustive global search ensures the general applicability of alignment. EMAlign is evaluated on 64 global map pairs, 195 local map pairs, and 60 structure-to-map pairs at 3–10 Å resolution and compared with gm-fit, fitmap, VESPER, and CryoAlign. It is shown that EMAlign outperforms the other methods in both global and local alignment, achieving mean RMSDs of 1.03 Å (global), 2.56 Å (local), and 0.82 Å (structure-to-map), with success rates of 100.0%, 100.0%, and 98.3% under the criterion of RMSD *<* 10 Å . The EMAlign package is freely available at https://github.com/huang-laboratory/EMAlign/.

## 1 Introduction

The rapid development of hardware^1^, sample preparation^2^, and image processing algorithms^3–6^ has established cryo-electron microscopy (cryo-EM) as a mainstream approach for determining the structures of biological macromolecules and their complexes^7–10^. With the expanding number of three-dimensional (3D) density maps deposited in the Electron Microscopy Data Bank (EMDB)^11^, cryo-EM provides an increasingly rich structural resource for studying biological function at the molecular level. However, fully exploiting this expanding archive requires computational tools that can reliably compare and superimpose related density maps.

Alignment of cryo-EM density maps is a fundamental step in many downstream analyses. Global alignment enables comparison of structures across functional states^12–14^, characterization of ligand-induced conformational changes^15^, and retrieval of related maps from large databases. Local alignment is essential when two maps share only partial structural overlap, as in subregion fitting, domain docking, and complex assembly^16, 17^. Accurate placement of atomic models within experimental density maps is likewise required for structure-to-map fitting and automated model building^18–21^. With increasing attention to conformational heterogeneity in single-particle cryo-EM reconstructions^22–25^, the demand for accurate, automatic, and broadly applicable map alignment has become increasingly urgent.

Several computational methods have been developed for cryo-EM map alignment. gmfit represents density maps as Gaussian mixture models (GMMs) and searches for the transformation that maximizes the correlation between GMMs^26^, providing an efficient reduced representation for global matching. fitmap, implemented in UCSF Chimera^27^, maximizes the local correlation between two maps starting from a user-provided initial placement and is widely used in interactive modeling workflows. More recently, VESPER^28^ and CryoAlign^29^ have achieved substantial improvements over earlier approaches. VESPER encodes each map as a set of vectors oriented toward local density maxima derived from mean-shift analysis and evaluates candidate alignments by the sum of dot products between matched vectors using FFT-based exhaustive search^30^. CryoAlign converts raw density maps into point clouds with local feature descriptors and performs alignment through coarse global registration followed by Sparse-ICP refinement^31, 32^. These methods have advanced global and local map alignment, map-to-map comparison, and database retrieval.

Despite present developments, accurate map alignment remains challenging. Methods that operate directly on raw voxel intensities are sensitive to low signal-to-noise ratios (SNR), incomplete sampling, and resolution variation across the map. fitmap depends on manual intervention and can fail when the starting pose is inaccurate. gmfit is affected by parameter choices in GMM representation and often trades accuracy for computational efficiency. Although VESPER captures local structural patterns effectively, its exhaustive search in rotational and translational space is computationally demanding. CryoAlign improves efficiency and accuracy relative to earlier methods, but its performance still depends on map resolution and, for local alignment, may benefit from auxiliary masking. A common limitation shared by these approaches is that alignment is attempted without first converting experimental maps into biologically focused, noise-suppressed representations.

To address these challenges, we present EMAlign, a fast Fourier transform (FFT)-based exhaustive search strategy for automatic global and local alignment of cryo-EM maps with predicted main-chain probability maps using deep learning. For map-to-map alignment, EMAlign converts each experimental input map into a main-chain probability map using BiMCUNet^33–35^, a bidirectional Mamba–convolutional network developed for cryo-EM map interpretation. Mean-shift clustering is then applied to extract local density-peak points (LDP), which are matched through FFT-based exhaustive search and simplex refinement in the EMAlign workflow. For structure-to-map fitting, BiMCUNet prediction is applied only to the target experimental map, whereas the moving atomic model is converted directly into LDP from backbone coordinates extracted from PDB/mmCIF files. By representing macromolecular backbones rather than raw voxels, EMAlign reduces sensitivity to noise while preserving the local structural features required for rigid-body superposition. With the FFT-based exhaustive search, EMAlign also does not rely on manual intervention and ensure the general applicability of alignment. EMAlign is evaluated on benchmark sets comprising 64 global map pairs, 195 local map pairs, and 60 structure-to-map pairs at 3–10 Å resolution and compared with gmfit, fitmap, VESPER, and CryoAlign. Our results demonstrate that EMAlign achieves improved alignment accuracy and success rates across the three benchmark tasks.

## 2 Results

### 2.1 Overview of EMAlign

Figure 1 summarizes the EMAlign pipeline and its two core algorithmic modules. Figure 1a shows the end-to-end workflow for cryo-EM map alignment. The input consists of a moving cryo-EM density map and a fixed reference map, which may contain proteins, nucleic acids, or their complexes. EMAlign first predicts main-chain and class probability maps for both experimental maps using a trained BiMCUNet network (Methods). Mean-shift clustering is then applied to extract categorized local density-peak points (LDP), in which protein and nucleic-acid peaks can be distinguished according to the predicted class map when class-map guidance is used. An FFT-based exhaustive match is performed on the LDP representations to enumerate candidate rigid-body transformations in rotation and translation space, followed by simplex refinement of the top-scoring solutions. The final transformation is used to superimpose the moving map and its associated atomic model onto the fixed map for downstream comparison and model building. Figure 1b details the BiMamba-Conv block in BiMCUNet, after a 1 *×* 1 convolution, the input features are split along the channel dimension and processed in parallel by a bidirectional BiMamba branch for global contextual features and a residual 3D convolutional branch with FRN normalization for local spatial features; the two branch outputs are concatenated and fused through a 1 *×* 1 convolution with a residual connection. Figure 1c illustrates the principle of FFT-based translational matching: voxelized LDP grids of the fixed and rotated moving maps are cross-correlated in the frequency domain to exhaustively evaluate translations for each sampled orientation; the highest-scoring candidates are then refined by simplex optimization.

**Fig. 1:**
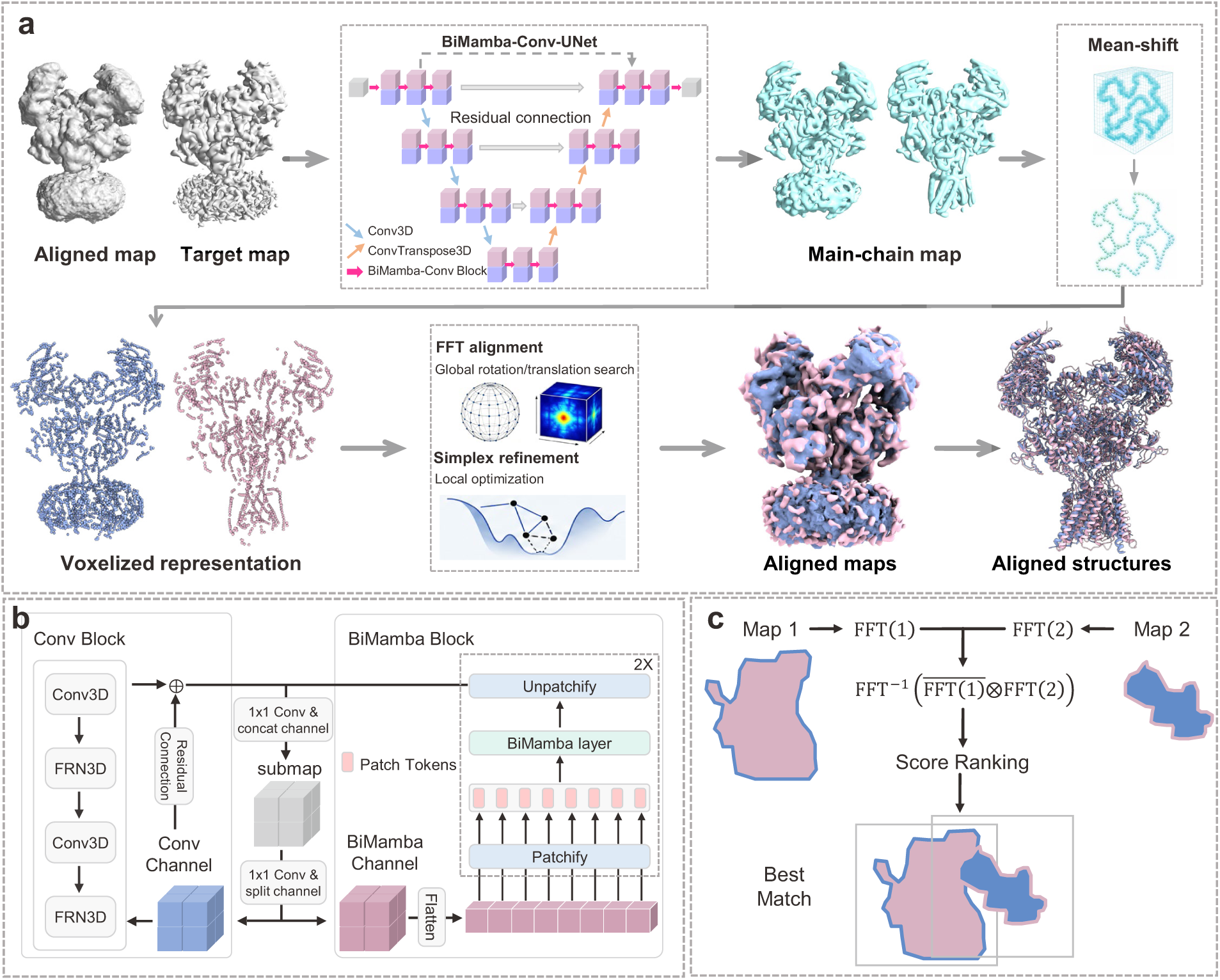
The workflow of EMAlign. **a**, Overall alignment pipeline. Two experimental cryo-EM maps (Aligned and Target) are converted into main-chain and class probability maps by BiMCUNet. Mean-shift clustering extracts local density-peak points (LDP), which are voxelized and matched by FFT-based exhaustive search over rotations and translations, followed by simplex refinement. The final rigid-body transform superimposes the moving map and optional atomic model onto the fixed map. **b**, Architecture of the BiMamba-Conv (BiMC) block in BiMCUNet. Input features are transformed by a 1 *×* 1 convolution and split along the channel dimension; one branch is processed by a bidirectional BiMamba block for global contextual features, and the other by a residual 3D convolutional block with FRN normalization for local spatial features; the two branch outputs are concatenated and fused through a 1 *×* 1 convolution with a residual connection. **c**, Principle of FFT-based translational matching. Voxelized main-chain grids of the fixed and rotated moving maps are transformed to the frequency domain (FFT), multiplied, and inverted (IFFT) to obtain a correlation-based score map; the translation with the best score is selected for each sampled orientation and ranked among candidates for simplex refinement.

To accelerate the alignment for large cryo-EM maps, EMAlign implements a GPU-accelerated alignment pipeline. BiMCUNet inference is performed on GPU, and the default Python backend offloads the most time-consuming steps, including mean-shift peak extraction, FFT-based translational search, and simplex refinement, to CUDA via Numba and CuPy. When CUDA is unavailable, the workflow automatically falls back to Fortran CPU implementations, preserving the same algorithmic steps while trading speed for hardware compatibility.

### 2.2 Performance of global alignment

We first evaluated EMAlign on a global alignment benchmark comprising 64 pairs of experimental cryo-EM maps. Fig. 2 shows the alignment results of EMAlign and four other methods including CryoAlign, VESPER, fitmap, and gmfit. It can be seen from the figure that EMAlign outperforms the other methods in both RMSD and success rates. Specifically, EMAlign achieves a mean RMSD of 1.03 Å over all 64 pairs with a 100.0% success rate. By comparison, CryoAlign yields a mean RMSD of 2.14 Å and an 87.5% success rate, VESPER shows a mean RMSD of 3.62 Å and a 73.4% success rate, fitmap obtains a mean RMSD of 1.28 Å but only a 50.0% success rate, and gmfit gives a mean RMSD of 3.40 Å and a 59.4% success rate. The box plots in Fig. 2a show that the RMSD distribution of EMAlign is markedly shifted toward lower values, with a median of 0.67 Å compared with 2.23 Å for CryoAlign and 4.26 Å for VESPER. Among all EMAlign results, 61% of the cases have RMSDs below 1 Å , indicating near-perfect superposition. Head-to-head comparisons in Fig. 2b further indicate that EMAlign generally achieves lower RMSDs than the other methods on a per-pair basis.

**Fig. 2:**
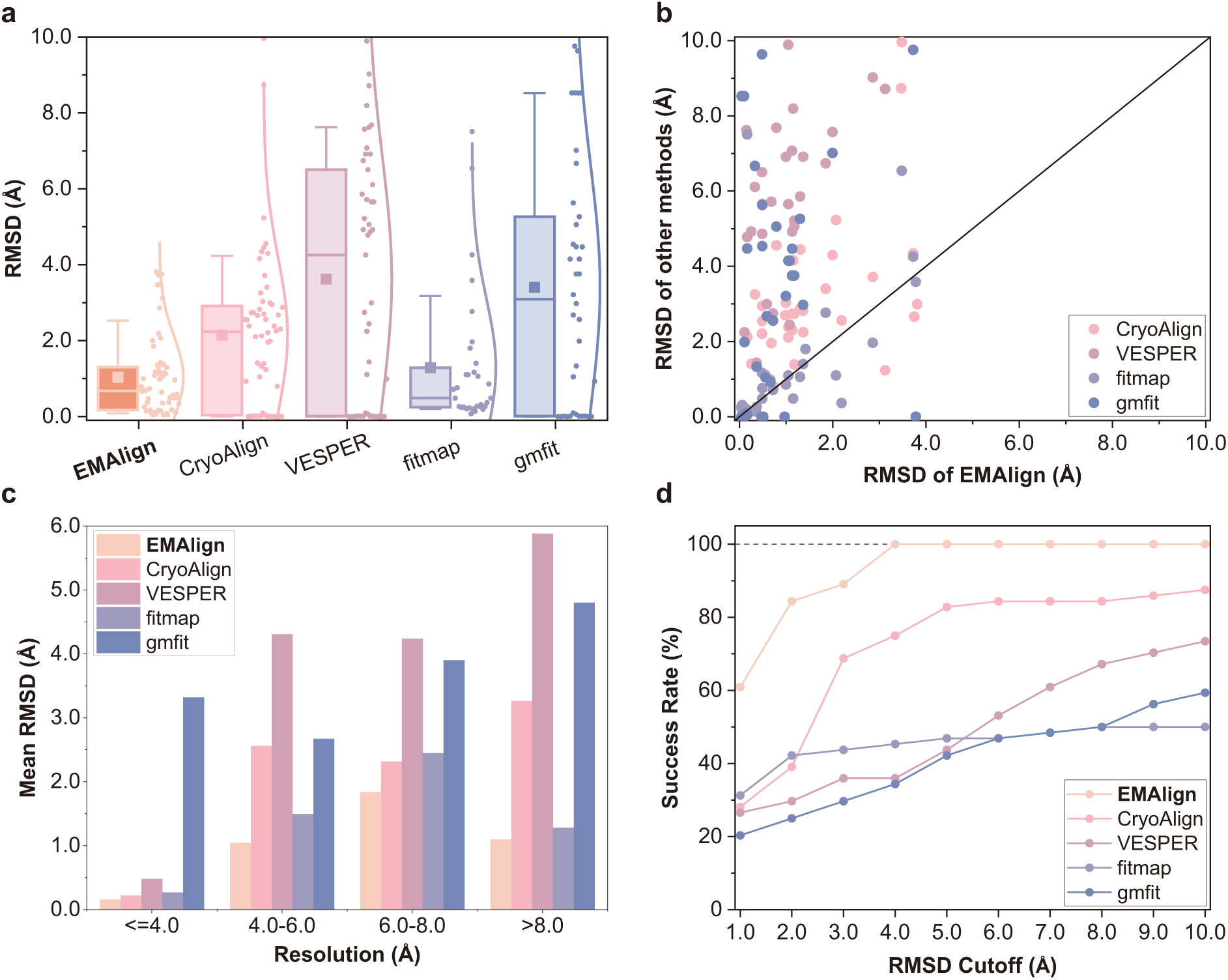
Evaluation of global alignment performance on. *n* **= 64 pairs of cryo-EM maps. a**, Box-and-whisker plots of RMSD for EMAlign, CryoAlign, VESPER, fitmap, and gmfit. The central line represents the median, with circles indicating the mean value. The lower and upper hinges correspond to the first and third quartiles, respectively, while the whiskers extend to 1.5 times the interquartile range. **b**, Head-to-head comparison of RMSD between EMAlign and the other methods for each map pair. **c**, Resolution-stratified mean RMSD. Resolution bins are defined by max(moving, fixed) resolution. **d**, Success rates at different RMSD cutoffs.

Resolution-stratified analysis (Fig. 2c and Table 1) demonstrates that EMAlign maintains high accuracy across the full resolution range. Specifically, EMAlign achieves 100.0% success in all four bins defined by max(moving resolution, fixed resolution) (*≤*4, 4–6, 6–8, and *>*8 Å), and yields the lowest mean RMSD in every bin. In the 4–6 Å bin, which contains the largest number of pairs (*n* = 27), EMAlign achieves a mean RMSD of 1.04 Å , compared with 2.56 Å for CryoAlign and 4.31 Å for VESPER. For more stringent RMSD cutoffs, the advantage of EMAlign becomes more pronounced (Fig. 2d). For example, at 4 Å , EMAlign succeeds for 100.0% of the cases, whereas CryoAlign and VESPER succeed for only 75.0% and 35.9% of the cases, respectively.

**Table 1:** Performance of global alignment by EMAlign, CryoAlign, VESPER, fitmap, and gmfit on 64 pairs of cryo-EM maps. The mean RMSD (Å) / success rate are reported for each reso-lution bin defined by max(moving, fixed) resolution. The numbers in bold fonts indicate the best performances for the corresponding categories.

| Resolution | **EMAlign** | CryoAlign | VESPER | fitmap | gmfit |
| --- | --- | --- | --- | --- | --- |
| $\leq 4$ Å ($n = 13$) | **0.15 / 100.0%** | 0.22 / 92.3% | 0.48 / 92.3% | 0.27 / 84.6% | 3.32 / 92.3% |
| 4–6 Å ($n = 27$) | **1.04 / 100.0%** | 2.56 / 77.8% | 4.31 / 66.7% | 1.49 / 33.3% | 2.67 / 51.9% |
| 6–8 Å ($n = 13$) | **1.84 / 100.0%** | 2.32 / 92.3% | 4.24 / 61.5% | 2.45 / 61.5% | 3.90 / 46.2% |
| $> 8$ Å ($n = 11$) | **1.09 / 100.0%** | 3.26 / 100.0% | 5.88 / 81.8% | 1.28 / 36.4% | 4.80 / 54.5% |
| All ($n = 64$) | **1.03 / 100.0%** | 2.14 / 87.5% | 3.62 / 73.4% | 1.28 / 50.0% | 3.40 / 59.4% |

Figure 3 shows two representative global alignment examples. In Fig. 3a, the alignment between the DNA-free and DNA-bound forms of the *E. coli* DNA polymerase complex (EMD-3201/PDB-5FKU and EMD-3202/PDB-5FKW) yields an RMSD of 0.49 Å for EMAlign, compared with 2.95 Å for CryoAlign, 4.86 Å for VESPER, and 4.54 Å for gmfit. In Fig. 3b, the alignment of the V-ATPase–SidK complex (EMD-8726/PDB-5VOZ) to the yeast V-ATPase state 3 (EMD-6286/PDB-3J9V) achieves an RMSD of 0.33 Å for EMAlign, whereas CryoAlign, VESPER, and gmfit yield the RMSDs of 3.26 Å , 6.11 Å , and 6.67 Å , respectively, and fitmap totally fails on this case. In both cases, the structures aligned by EMAlign closely match the MM-align references, whereas the other methods show noticeable misplacement.

**Fig. 3:**
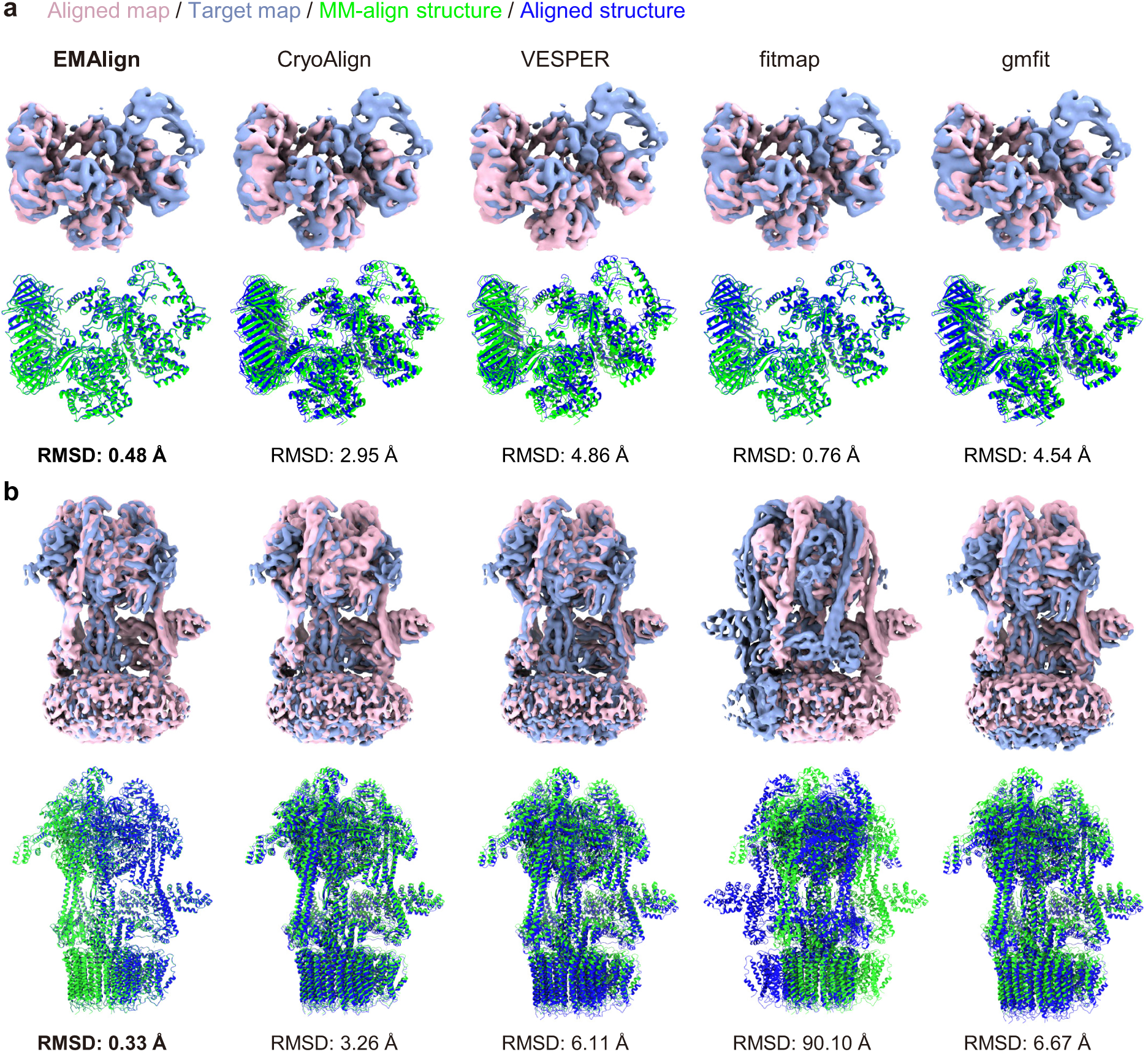
Examples of global alignment results. The target maps are colored in purple, the aligned maps are in light blue, the MM-align–superimposed structures are in green, and the aligned structures are in blue. **a**, Global alignment of EMD-3201 (PDB ID: 5FKU) at 8.34 Å resolution to EMD-3202 (PDB ID: 5FKW) at 7.3 Å resolution. **b**, Global alignment of EMD-8726 (PDB ID: 5VOZ) at 7.6 Å resolution to EMD-6286 (PDB ID: 3J9V) at 8.3 Å resolution.

### 2.3 Performance of local alignment

Global alignments are appropriate when two maps are of comparable size and share extensive structural overlap. In practice, however, one map often represents only a subregion of another, or two maps share only partial overlap. We therefore evaluated EMAlign on a local alignment benchmark comprising 195 map pairs at 3–10 Å resolution. Fig. 4 shows the corresponding alignment results of EMAlign and the other four methods including CryoAlign, VESPER, fitmap, and gmfit, again demonstrating the best performance of EMAlign.

**Fig. 4:**
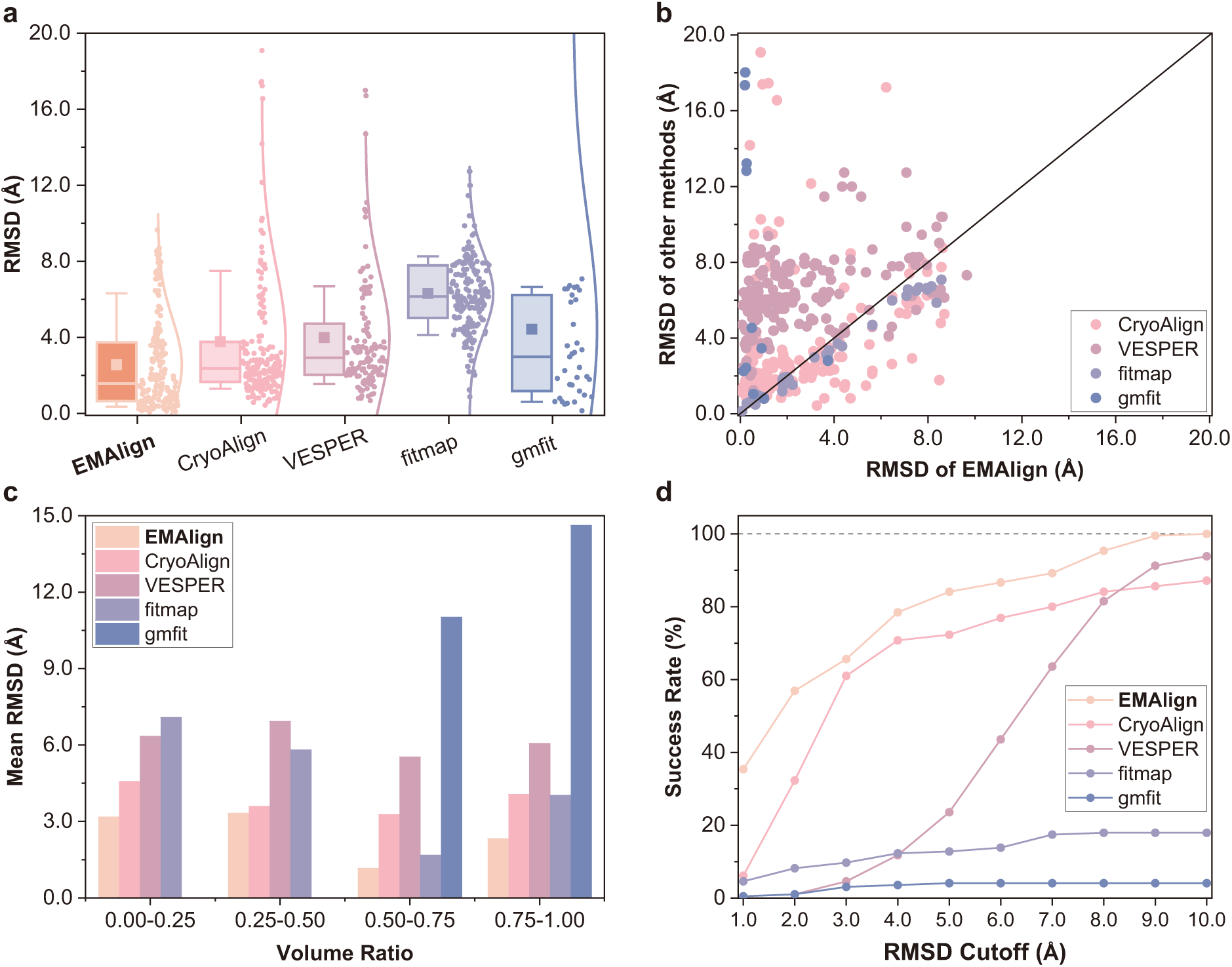
Evaluation of local alignment performance on. *n* **= 195 pairs of cryo-EM maps. a**, Box-and-whisker plots of RMSD for EMAlign, CryoAlign, VESPER, fitmap and gmfit. The central line represents the median, with circles indicating the mean value. The lower and upper hinges correspond to the first and third quartiles, respectively, while the whiskers extend to 1.5 times the interquartile range. **b**, Head-to-head comparison of RMSD between EMAlign and the other methods for each map pair. **c**, Volume-ratio-stratified mean RMSD. Volume ratio bins are defined by source-to-target map volume ratio (moving map volume divided by fixed map volume). **d**, Success rates at different RMSD cutoffs.

As shown in Fig. 4a, EMAlign achieves a mean RMSD of 2.56 Å over all 195 pairs with a 100.0% success rate at the 10 Å cutoff. CryoAlign yields a mean RMSD of 3.78 Å and an 87.2% success rate, VESPER gives a mean RMSD of 6.32 Å and a 93.8% success rate, fitmap obtains a mean RMSD of 4.43 Å and a 17.9% success rate, and gmfit shows a mean RMSD of 13.49 Å and a 4.1% success rate.

Although VESPER obtains valid results for most of the cases, its median RMSD of 6.16 Å indicates limited precision for high-accuracy local fitting. By contrast, the RMSD distribution of EMAlign is concentrated at substantially lower values, with a median of 1.58 Å . Head-to-head comparisons in Fig. 4b show that EMAlign yields lower RMSD than CryoAlign and VESPER for 108 and 172 of the valid pairwise comparisons, respectively.

Volume-ratio-stratified analysis (Fig. 4c and Table 2) shows that EMAlign remains accurate even when the moving map occupies only a small fraction of the fixed map. Specifically, EMAlign achieves 100.0% success in all four bins defined by source-to-target map volume ratio and delivered the lowest mean RMSD in every bin. In the smallest-ratio bin (0–0.25; *n* = 18), where the moving map occupies at most one quarter of the fixed map volume, EMAlign achieves a mean RMSD of 3.17 Å , compared with 4.57 Å for CryoAlign and 6.34 Å for VESPER. In the 0.5–0.75 bin (*n* = 38), EMAlign achieves a mean RMSD of 1.16 Å with 100.0% success. At the 10 Å success cutoff (Fig. 4d), EMAlign succeeds in 100.0% of pairs, compared with 87.2% for CryoAlign and 93.8% for VESPER.

**Table 2:** Performance of local alignment by EMAlign, CryoAlign, VESPER, fitmap, and gmfit on 195 pairs of cryo-EM maps. The mean RMSD (Å) / success rate are reported by source-to-target map volume ratio (moving map volume divided by fixed map volume). The numbers in bold fonts indicate the best performances for the corresponding categories. “–” means that a method totally fails for the category.

| Volume ratio | **EMAlign** | CryoAlign | VESPER | fitmap | gmfit |
| --- | --- | --- | --- | --- | --- |
| 0–0.25 ($n = 18$) | **3.17 / 100.0%** | 4.57 / 88.9% | 6.34 / 83.3% | 7.08 / 5.6% | – / 0.0% |
| 0.25–0.5 ($n = 75$) | **3.33 / 100.0%** | 3.60 / 93.3% | 6.92 / 92.0% | 5.81 / 14.7% | – / 0.0% |
| 0.5–0.75 ($n = 38$) | **1.16 / 100.0%** | 3.27 / 78.9% | 5.53 / 97.4% | 1.68 / 10.5% | 11.02 / 5.3% |
| 0.75–1 ($n = 64$) | **2.33 / 100.0%** | 4.06 / 84.4% | 6.07 / 96.9% | 4.03 / 29.7% | 14.62 / 9.4% |
| All ($n = 195$) | **2.56 / 100.0%** | 3.78 / 87.2% | 6.32 / 93.8% | 4.43 / 17.9% | 13.49 / 4.1% |

Two illustrative local alignment examples are shown in Fig. 5. In Fig. 5a, the alignment of the *Saccharomyces cerevisiae* 80S ribosome bound to eEF2 and a viral IRES (EMD-6647/PDB-5JUU) to the porcine mitochondrial 39S large subunit (EMD-2490/PDB-4CE4) yields an RMSD of 1.49 Å for EMAlign, whereas VESPER produces an RMSD of 7.74 Å and CryoAlign fails to give a valid result. In Fig. 5b, the alignment of the full human mitochondrial ribosome (EMD-2876/PDB-3J9M) to the human mitochondrial large subunit (EMD-2762/PDB-3J7Y) shows an RMSD of 0.25 Å for EMAlign, compared with 2.42 Å for CryoAlign, 2.48 Å for VESPER, and 2.42 Å for gmfit. For both examples, EMAlign preserves the local structural correspondence visible in the MM-align references, whereas the other methods show clear misplacement of the moving substructure.

**Fig. 5:**
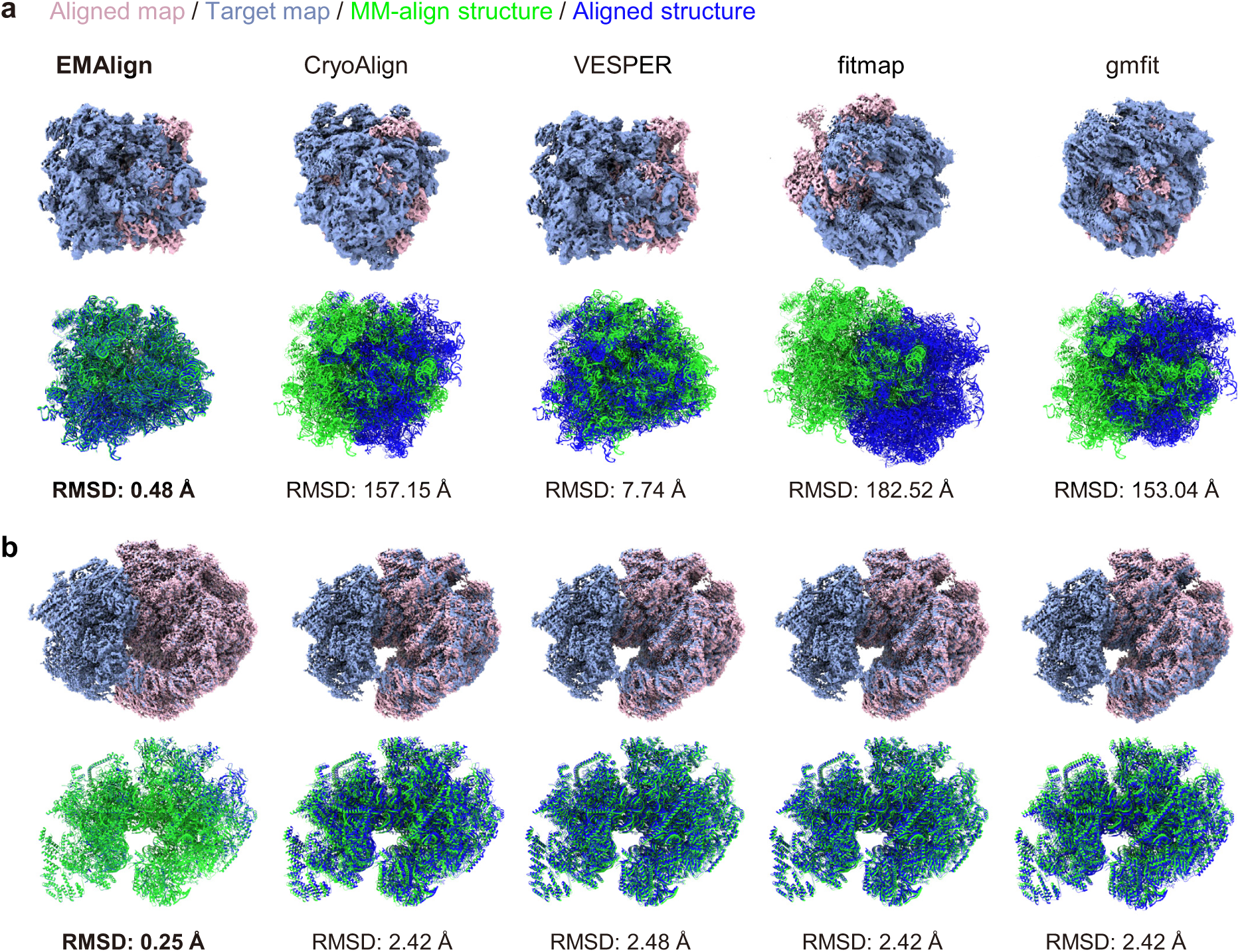
Examples of local alignment results. The target maps are colored in purple, the aligned maps are in light blue, the MM-align–superimposed structures are in green, and the aligned structures are in blue. **a**, Local alignment of EMD-6647 (PDB ID: 5JUU) at 4.0 Å resolution to EMD-2490 (PDB ID: 4CE4) at 4.9 Å resolution. **b**, Local alignment of EMD-2876 (PDB ID: 3J9M) at 3.5 Å resolution to EMD-2762 (PDB ID: 3J7Y) at 3.4 Å resolution.

### 2.4 Structure-to-map alignment

Besides map-to-map alignment, EMAlign also supports structure-to-map fitting, in which an atomic model is aligned into an experimental cryo-EM density map. Therefore, we compared EMAlign with CryoAlign and VESPER on a benchmark of 60 structure-to-map pairs derived from the global evaluation set after removing redundant entries. During the evaluations, CryoAlign and VESPER generate simulated density maps from atomic models and align them to the target map. EMAlign adopts an asymmetric representation strategy: the target experimental map is processed by BiMCUNet and converted into LDP via mean-shift clustering, whereas the moving atomic model bypasses neural network prediction entirely. Backbone heavy-atom coordinates are extracted directly from the PDB/mmCIF file and downsampled into structure-derived LDP anchor points for alignment evaluations.

Fig. 6 shows the structure-to-map alignment of EMAlign and two other methods including CryoAlign and VESPER. It can be seen from the figure that EMAlign performs the best among the compared methods. Specifically, EMAlign achieves a mean RMSD of 0.82 Å with a 98.3% success rate over all 60 pairs (one failure). By comparison, CryoAlign yields a mean RMSD of 2.60 Å with a 60.0% success rate, and VESPER obtains a mean RMSD of 1.59 Å with a 78.3% success rate. The box plots in Fig. 6a show that EMAlign consistently produces the lowest RMSD values, with a median of 0.43 Å compared with 2.58 Å for CryoAlign and 1.37 Å for VESPER. Head-to-head comparisons in Fig. 6b show that EMAlign yields lower RMSD than CryoAlign and VESPER in 30 of 36 and 45 of 47 valid pairwise comparisons, respectively.

**Fig. 6:**
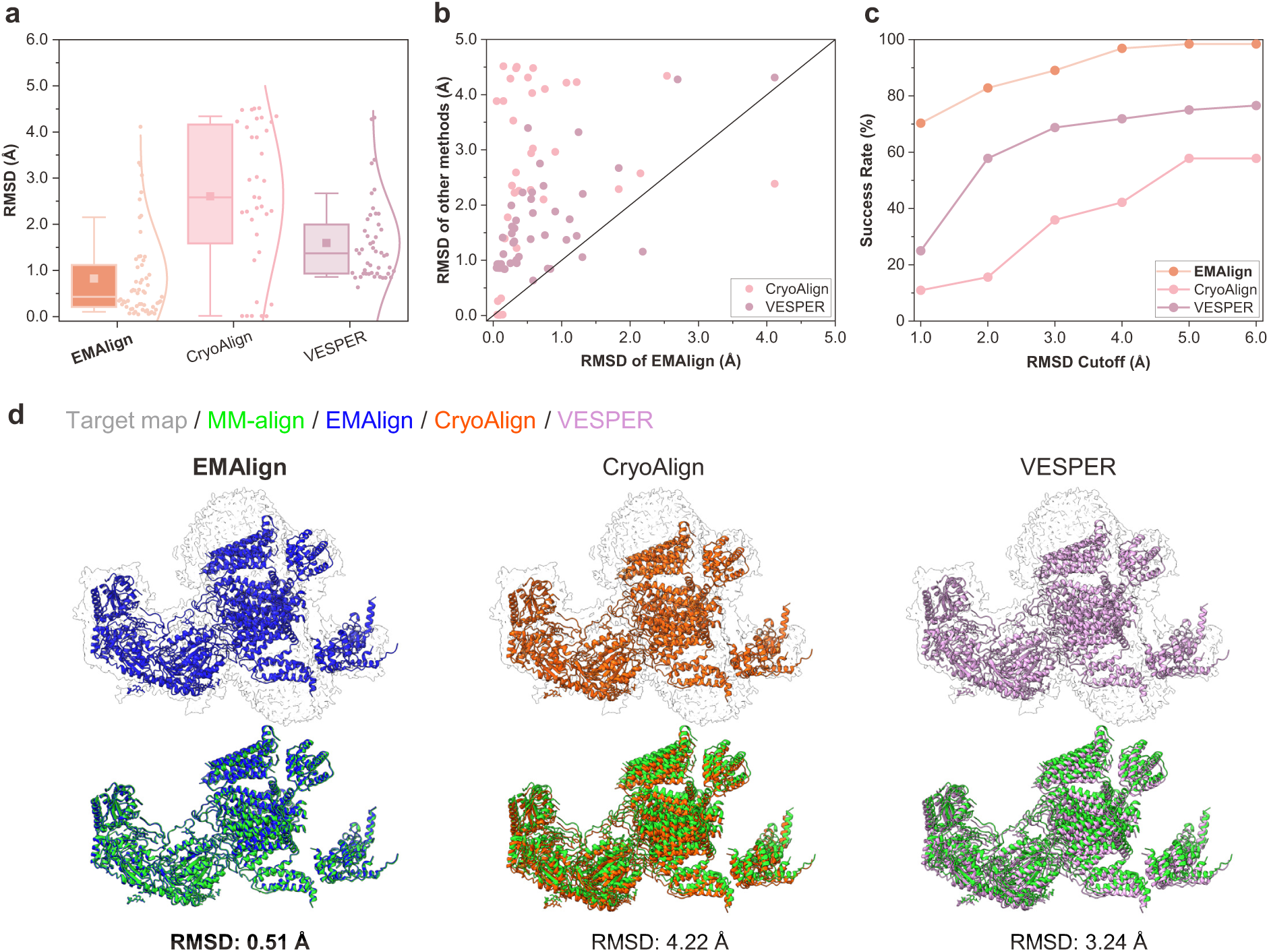
Evaluation of structure-to-map alignment performance on. *n* **= 60 pairs of structures and maps. a**, Box-and-whisker plots of RMSD for EMAlign, CryoAlign and VESPER. The central line represents the median, with circles indicating the mean value. The lower and upper hinges correspond to the first and third quartiles, respectively, while the whiskers extend to 1.5 times the interquartile range. **b**, Head-to-head comparison of RMSD between EMAlign and the other methods for each pair. **c**, Success rates at different RMSD cutoffs. **d**, Structure-to-map fitting of PDB-3JBR at 6.1 Å resolution into EMD-9515 (PDB ID: 5GJW) at 3.9 Å resolution by EMAlign, CryoAlign and VESPER. The MM-align reference is in green, the EMAlign-fitted model is in blue, the CryoAlign-fitted model is in orange, and the VESPER-fitted model is in pink.

Resolution-stratified analysis (Table 3) demonstrates robust performance across resolution bins defined by max(structure resolution, target map resolution). In the *≤*4 Å bin (*n* = 13), EMAlign achieved a mean RMSD of 0.17 Å and 100.0% success. In the 4–6 Å bin (*n* = 24), EMAlign achieves a mean RMSD of 0.84 Å and a 95.8% success rate, whereas CryoAlign succeeds for only 45.8% of the cases with a mean RMSD of 3.39 Å . At the 10 Å success cutoff, EMAlign maintains high coverage across all bins (Fig. 6c), substantially outperforming CryoAlign and VESPER in the mid-resolution range where structure-to-map fitting is most challenging.

**Table 3:** Performance of structure-to-map alignment accuracy of EMAlign, CryoAlign, and VESPER on 60 pairs of structures and maps. The mean RMSD (Å) / success rate are reported for each resolution bin defined by max(structure, target map) resolution. The numbers in bold fonts indicate the best performances for the corresponding categories.

| Resolution | EMAlign | CryoAlign | VESPER |
| --- | --- | --- | --- |
| $\leq 4 \text{ \AA}$ ($n = 13$) | **0.17 / 100.0%** | 0.55 / 69.2% | 1.05 / 84.6% |
| 4–6 $\text{\AA}$ ($n = 24$) | **0.84 / 95.8%** | 3.39 / 45.8% | 1.39 / 75.0% |
| 6–8 $\text{\AA}$ ($n = 12$) | **1.32 / 100.0%** | 3.13 / 75.0% | 1.91 / 66.7% |
| $> 8 \text{ \AA}$ ($n = 11$) | **1.02 / 100.0%** | 3.32 / 63.6% | 2.29 / 90.9% |
| All ($n = 60$) | **0.82 / 98.3%** | 2.60 / 60.0% | 1.59 / 78.3% |

An illustrative example is shown in Fig. 6d for fitting the structure of the rabbit voltage-gated calcium channel Ca_v_1.1 complex (PDB-3JBR) into its corresponding cryo-EM density map (EMD-9515, targeted by PDB-5GJW). On this case, EMAlign achieves an RMSD of 0.51 Å , whereas CryoAlign and VESPER yield the RMSDs of 4.22 Å and 3.24 Å , respectively. The model fitted by EMAlign closely matches the MM-align reference, whereas the CryoAlign- and VESPER-fitted models show noticeable displacement from the target density.

## 3 Discussion

In this study, we present EMAlign, a deep-learning-based method for automatic global and local alignment of cryo-EM maps. EMAlign converts experimental raw maps into biologically interpretable local density-peak points (LDP) through BiMCUNet-based prediction and mean-shift clustering. The accurate alignment is performed using FFT-based exhaustive match and simplex refinement. In addition, EMAlign can also be used for structure-to-map fitting, where the moving model is represented directly by backbone-derived LDP without neural network inference on the atomic coordinates. A GPU-accelerated implementation enables efficient processing of large maps. By representing macro-molecular backbones as compact LDP rather than raw voxel intensities, EMAlign reduces sensitivity to noise while preserving the local structural features required for accurate rigid-body matching.

EMAlign was benchmarked on 64 global map pairs, 195 local map pairs, and 60 structure-to-map pairs at 3–10 Å resolution and compared with gmfit, fitmap, VESPER, and CryoAlign. Across these tests, EMAlign achieves the lowest overall RMSD and the highest success rates under a unified 10 Å success cutoff for all three benchmark tasks. The corresponding mean RMSDs are 1.03 Å (global), 2.56 Å (local), and 0.82 Å (structure-to-map), with success rates of 100.0%, 100.0%, and 98.3%, respectively. EMAlign also remains accurate across different resolution and volume-ratio ranges and in representative examples spanning diverse complexes. These results indicate that EMAlign is well suited for comparing maps across functional states, searching map libraries for related structures, and providing initial structure placements for atomic model building.

With the rapid growth of cryo-EM and cryo-ET depositions, reliable map alignment is becoming increasingly important for mining structural databases, characterizing conformational landscapes, and linking experimental density to computational models. Future development of EMAlign may include tighter integration with map post-processing and automated model-building workflows, extension to lower-resolution and tomography maps, and improved handling of compositional and conformational heterogeneity beyond rigid-body superposition. Class-aware matching for protein–nucleic acid assemblies is available as an optional workflow, and tighter integration with emerging generative structure-prediction models represents an additional promising direction. It is anticipated that EMAlign will serve as a practical and accurate component in automated cryo-EM structure-determination pipelines.

## 4 Methods

### 4.1 Training of main-chain prediction network

EMAlign employs the BiMCUNet^33–35^ for main-chain probability prediction from raw cryo-EM maps, and performs map alignment through a fast Fourier transform (FFT)-based exhaustive match. Rather than aligning raw experimental maps directly, EMAlign first converts each experimental input map into a main-chain probability map and a residue-class probability map using this network. The main-chain map estimates the likelihood that a voxel lies near a macromolecular backbone atom, whereas the class map assigns each voxel to background, protein backbone, or nucleic-acid backbone. Compared with raw density maps, these predictions suppress noise and emphasize biologically interpretable features, thereby improving subsequent alignment accuracy.

The BiMCUNet is a bidirectional Mamba–convolutional UNet built upon selective state space models^34, 35^. As shown in Fig. 1b, each BiMamba-Conv block first applies a 1 *×* 1 convolution and splits the features along the channel dimension into two equal partitions, which are processed in parallel by a BiMamba branch and a residual 3D convolutional branch with FRN normalization, respectively; the branch outputs are concatenated and fused by a 1 *×* 1 convolution with a residual connection to combine global and local features. Training and validation are based on the 348 and 65 curated EMDB maps at 2–10 Å from the EMReady2 study^33^, respectively, which include protein-only, nucleic-acid-containing, and low-resolution cryo-EM maps. For each entry, backbone heavy atoms were extracted from the deposited atomic model: N, C*α*, and C for proteins, and P, O5^′^, C5^′^, C4^′^, C3^′^, and O3^′^ for nucleic acids. The main-chain target map is simulated from these coordinates with UCSF Chimera^27^ `molmap` at the reported map resolution. A voxel-wise class label map is derived from residue type, with labels of 0 (background), 1 (protein), and 2 (nucleic acid).

All maps are resampled to 1.0 Å voxel spacing by cubic interpolation. Experimental maps are clipped at the 99.99th percentile of positive densities, scaled to the unit interval, and z-score normalized. Overlapping subvolumes of size 64 *×* 64 *×* 64 with a stride of 32 are extracted from each aligned experimental, main-chain, and class map triplet; subvolumes with fewer than 5% foreground (non-background) voxels are discarded. During training, BiMCUNet simultaneously regresses the main-chain probability map and classifies each voxel into the three residue categories. Given a predicted main-chain subvolume **Y**^pred^ and its ground-truth target **Y**^GT^ of shape *M ×M ×M* (*M* = 64), the smooth *L*_1_ loss is defined as

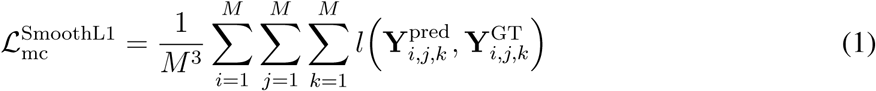

where 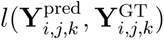 is the smooth *L*_1_ distance at position (*i, j, k*):

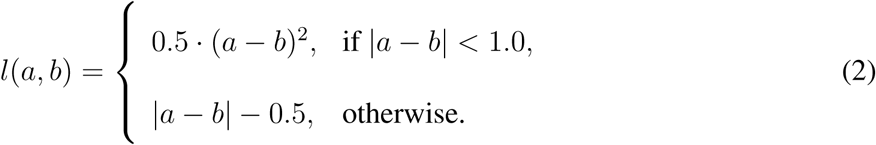

The SSIM loss measures structural similarity between the predicted and target main-chain subvolumes^36^:

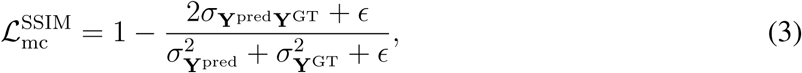

where *σ*_Y_pred and *σ*_Y_GT are the standard deviations, *σ*_Y_pred_Y_GT is the covariance, and *ɛ* = 10^−6^. For class prediction, weighted cross-entropy and multiclass Dice losses are applied to the three-class label map, with higher weights assigned to protein and nucleic-acid categories to mitigate class imbalance. The total training objective is defined by the following loss function.

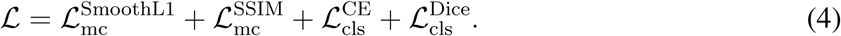

The model is optimized with AdamW (learning rate 10^−4^) for 200 epochs using mixed-precision training; random rotational augmentation is applied to input subvolumes. The checkpoint with the lowest validation loss is retained for inference. At runtime, experimental maps are processed with the same 1.0 Å interval resampling and tiled inference scheme (64^3^ boxes, stride 32) to generate main-chain and class probability maps.

### 4.2 Main-chain representation of cryo-EM maps

Each map to be aligned is represented by a set of local density-peak points (LDP). For map-to-map alignment, both moving and fixed experimental maps are processed by BiMCUNet before LDP extraction. For structure-to-map fitting, BiMCUNet prediction is applied only to the target experimental map; the moving atomic model is not passed through the neural network. Instead, backbone heavy-atom coordinates are extracted directly from the PDB/mmCIF file using the same atom sets as in network training, and representative LDP anchor points are generated from these coordinates without simulated map generation.

After obtaining the predicted main-chain and class maps (or structure-derived coordinates), EMAlign converts each representation into local density-peak points (LDP) using a mean-shift algorithm for experimental maps, or direct coordinate sampling for atomic models. The grid points above a probability threshold are iteratively shifted toward local maxima of the main-chain map:

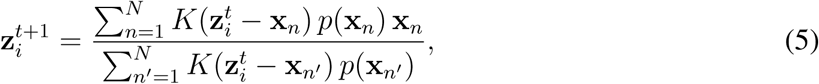

where **x***_n_* is a grid point, *p*(**x***_n_*) is the corresponding main-chain probability, and *K*(**r**) = exp(−*k*||*r*||^2^) is a Gaussian kernel with resolution-dependent parameter *k*. Shifted points are merged into LDP when their separation is below 2.0 Å ; the highest-probability point in each cluster is retained. Each LDP carries a main-chain density score and a categorical label. When class-map guidance is used, labels are assigned by majority voting over the predicted class map within a 3.0 Å radius around each LDP; protein and nucleic-acid peaks are labeled as 1 and 2, respectively. For structure-derived LDP, labels are assigned directly from residue type using the same backbone atoms as in network training. Users may choose whether to apply class-map guidance during mean-shift and FFT matching depending on the composition of the maps or models being aligned.

### 4.3 Map alignment through an FFT-based exhaustive match

Figure 1c schematically depicts the FFT-based translational search. For each sampled orientation of the moving map, the fixed and rotated moving LDP grids are transformed to the frequency domain, multiplied element-wise, and inverted to obtain a correlation-based score map; the translation with the optimal score is selected and the top candidates are retained for subsequent refinement. EMAlign next performs an FFT-based exhaustive search on the LDP representations of the moving (or source) and fixed (or target) maps to identify the optimal rigid-body transformation, followed by simplex refinement of the top-scoring candidates. Each LDP set is voxelized into a cubic grid of size *M × M ×M* with 3.0 Å spacing^37^. Rotational space is sampled at 15° intervals in Euler-angle space (4,392 orientations). For each orientation of the moving map, translational offsets are evaluated exhaustively by FFT cross-correlation. Let *X_l,m,n_* and *Y_l,m,n_* denote the fixed and rotated moving main-chain density grids, respectively, voxelized from the two LDP sets on cubic grids of size *M × M × M* . Let *λ* denote the average resolution of the two maps, *θ* = (*λ/π*)^1.5^ the resolution-dependent scaling factor, and 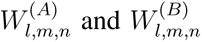 the optional class-weight factors assigned to voxels on the fixed and moving grids, respectively. The FFT match score for translation (*i, j, k*) is

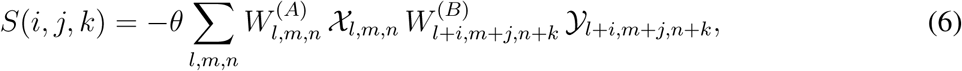

where a lower score indicates a better match. When class-map guidance is not used, 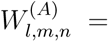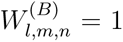 for all voxels and Eq. 6 reduces to a standard main-chain cross-correlation. When class-map guidance is used, each voxel inherits the label of the contributing LDP peak, and the weights are set to +1 for protein, *−*1 for nucleic acid, and 0 for background. The orientation–translation pair with the optimal score is selected, and the highest-ranked FFT solutions are refined on the LDP coordinates using simplex optimization with 1.5 Å grid spacing.

### 4.4 Evaluation of map alignment methods

EMAlign was extensively evaluated on three benchmark sets of 64 global map pairs, 195 local map pairs, and 60 structure-to-map pairs from the CryoAlign study^29^. For comparison, the alignments of four other methods, including fitmap, gmfit, VESPER, and CryoAlign were also computed or taken from the literature^29^. Default parameters were used for the compared methods. fitmap and gmfit were evaluated only on the map-to-map tasks. For structure-to-map fitting, EMAlign was compared with CryoAlign and VESPER. The alignment accuracy between the target map and the aligned map is measured by the root mean square deviation (RMSD) between the structure of the aligned map after map alignment and the structure of the aligned map after superimposition onto the structure of the target map with the MM-align program ^38^. An alignment was considered successful if the RMSD is below 10 Å , unless otherwise specified.

## Data availability

All published data sets used in this paper were taken from the EMDB and PDB. Source data are provided with this paper.

## Code availability

The EMAlign package is freely available for academic or non-commercial users at https://github.com/huang-laboratory/EMAlign or http://huanglab.phys.hust.edu.cn/EMAlign/.

## Acknowledgements

This work was supported by the National Natural Science Foundation of China (grants No. 32430020, 32161133002, and 62072199 to S.H.), the Postdoctoral Fellowship Program of CPSF (grant No. GZB20250617 to T.L) and the startup grant of Huazhong University of Science and Technology (to S.H.).

## Author contributions

S.H. conceived and supervised the project. H.C., J.C. and S.H. designed and performed the experiments. All authors analyzed the data. H.C., J.C. and S.H. wrote the paper. All authors read and approved the final version of the paper.

